# Optimizing genomics pipeline execution with integer linear programming

**DOI:** 10.1101/2024.02.06.579197

**Authors:** Olesya Melnichenko, Venkat S. Malladi

## Abstract

In the field of genomics, bioinformatics pipelines play a crucial role in processing and analyzing vast biological datasets. These pipelines, consisting of interconnected tasks, can be optimized for efficiency and scalability by leveraging cloud platforms such as Microsoft Azure. The choice of compute resources introduces a trade-off between cost and time. This paper introduces an approach that uses Linear Programming (LP) to optimize pipeline execution. We consider optimizing two competing cases: minimizing cost with a run duration restriction and minimizing duration with a cost restriction. Our results showcase the utility of using LP in guiding researchers to make informed compute decisions based on specific data sets, cost and time requirements, and resource constraints.

## 1 Introduction

In the realm of genomics, bioinformatics pipelines are integral to processing and analyzing vast amounts of biological data. A pipeline, in this context, is a series of data processing steps, or tasks, that are interconnected. Each task takes a set of inputs, processes it, and produces a set of outputs that may be used as inputs for the subsequent tasks or are considered terminal outputs. This sequence of tasks can be distributed, or divided into smaller, more manageable pieces or shards, to enhance efficiency and scalability. These pipelines can be run on cloud platforms like Microsoft Azure, leveraging its robust computational resources and storage capabilities. These pipelines are typically run using some of the following workflow managers all able to run on Azure: Nextflow [1, 2], Cromwell [3, 4], Snakemake [5, 6].

It’s important to note that the cost and duration of running these pipelines on Azure are influenced by the choice of compute resources. More powerful resources can process data faster but at a higher cost, necessitating a balance between speed, efficiency, and budget. Choosing the appropriate compute resources is dependent on the tools and data storage (memory and disk) requirements of each task.

With a variety of compute resources available, there is always a trade-off between cost and time when running a genomics pipeline. The preference for this trade-off depends on the researcher’s circumstances. If the researcher has limited funding and is less restricted on time, they might be interested in getting results as cheaply as possible without spending more than *T*_*max*_ hours per sample. In clinical settings where time is critical, the preference might be to get results as quickly as possible.

Efficient resources utilization remains a key issue in parallel and distributed computing environments. Resource allocation and task scheduling problems are well-known in this field, and a lot of effort has been invested in improving the management of Cloud resources [7, 8, 9, 10]. In particular, cost optimization of scientific workflows has been a focus of attention [11]. Existing open-source workflow engines in genomics field, such as CromwellOnAzure [12], Snakemake [5] or Nextflow [1], require researchers to provide a virtual machine-to-task assignment list in advance as one of the workflow inputs or make a default assignment with no guarantees for optimal performance.

When a researcher’s institution does not use a workflow management system that monitors workflow execution and optimizes task scheduling, the researcher faces the challenge of optimizing a pipeline according to project requirements while taking into account the variety of compute options and non-linear pipeline topology. In this paper, we demonstrate that linear programming can be a useful tool for pipeline execution optimization in a bioinformatics researcher’s toolbox. We consider **two use cases** for pipeline optimization:

**UC 1** minimizing the cost of running the pipeline with a restriction on run duration,

**UC 2** minimizing the duration of the pipeline run with a restriction on cost.

Genomics workloads often involve running a given pipeline on a large number of samples. Typically, data from these samples have the same number of reads, length, coverage and quality level, i.e. the sample dataset is homogeneous. Given that the dataset is homogeneous, we assume that average execution statistics collected for a subset of data are sufficient for preliminary pipeline performance estimates and optimizations. We also assume that statistics have been collected, and the scope of this work includes the following steps performed by the researcher: selecting a sample subset, selecting a virtual machine subset for each pipeline task, executing the pipeline on the selected subset of samples and virtual machines, collecting execution statistics, and computing the average cost and execution time for each pipeline task. For example, if virtual machine assignment is described by Table 1, then corresponding execution statistics will define matrices *C* = {*c*_*ij*_} and *T* = {*t*_*ij*_} presented as Tables 2 and 3.

**Table 1:** Set of experiments defined for a given pipeline.

|  | *Experiment<sub>1</sub>* | *Experiment<sub>2</sub>* |
| --- | --- | --- |
| *Task<sub>1</sub>* | $VM_{11}$ | $VM_{12}$ |
| *Task<sub>2</sub>* | $VM_{21}$ | $VM_{22}$ |
| *Task<sub>3</sub>* | $VM_{31}$ | $VM_{32}$ |

**Table 2:** Expected execution statistics: average time to complete each pipeline task.

|  | *Experiment<sub>1</sub>* | *Experiment<sub>2</sub>* |
| --- | --- | --- |
| *Task<sub>1</sub>* | $t_{11}$ | $t_{12}$ |
| *Task<sub>2</sub>* | $t_{21}$ | $t_{22}$ |
| *Task<sub>3</sub>* | $t_{31}$ | $t_{32}$ |

**Table 3:** Expected execution statistics: average cost to complete each pipeline task.

|  | *Experiment<sub>1</sub>* | *Experiment<sub>2</sub>* |
| --- | --- | --- |
| *Task<sub>1</sub>* | $c_{11}$ | $c_{12}$ |
| *Task<sub>2</sub>* | $c_{21}$ | $c_{22}$ |
| *Task<sub>3</sub>* | $c_{31}$ | $c_{32}$ |

## 2 Formulating linear programming problems

Maximizing or minimizing a function subject to constraints is a common problem in many fields. Linear programming (LP) is a special case of mathematical optimization that deals with linear functions and constraints. In LP, we assume that the function to be maximized or minimized is a linear function and that the constraints are linear equations or linear inequalities [13]. LP has proven useful in modeling diverse types of problems in planning, routing, scheduling, assignment, and design. In this section, we formulate LP problems for the use cases defined above.

Suppose we have a pipeline consisting of *n* tasks and *m* experiments defined by the researcher. The cost and time to complete each task are given by matrices *C* = {*c*_*ij*_} and *T* = {*t*_*ij*_}, respectively, where *i* = 1, …, *n* and *j* = 1, …, *m*. We can formulate the following multiple-choice problems for the use cases in consideration:

**UC 1** For each task, pick exactly one virtual machine out of *m* options in order to minimize the total cost of running the pipeline without exceeding the allotted time *T*_*max*_.

**UC 2** For each task, pick exactly one virtual machine out of *m* options in order to minimize the duration of the pipeline run without exceeding the allowed cost *C*_*max*_.

We can formulate LP problems for pipeline optimization by following the approach used for the multiple-choice knapsack problem (MCKP) [14]. To do this, we introduce indicator variables *I*_*ij*_, where *i* = 1, …, *n, j* = 1, …, *m*. The variable *I*_*ij*_ is equal to 1 if we choose virtual machine (VM) from *Experiment*_*j*_ for *Task*_*i*_, and 0 otherwise.

### 2.1 Linear pipeline topology

A linear pipeline consists of a chain of tasks arranged so that the output of each element is the input of the next. The processing of data is done in a linear and sequential manner, see example on Figure 1. In the case of a linear topology, the total time and cost to run a pipeline can be computed as a sum of all tasks time and cost. Following MCKP approach, the linear programming formulations can be expressed as:

**Figure 1:**
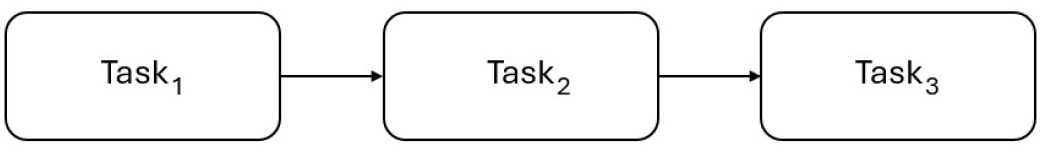
Example of linear pipeline topology

**LP 1 – Minimize cost subject to time constraint**

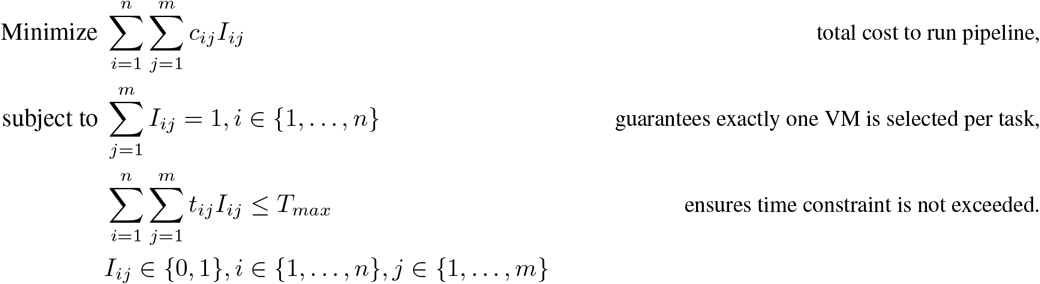

**LP 2 – Minimize time subject to cost constraint**

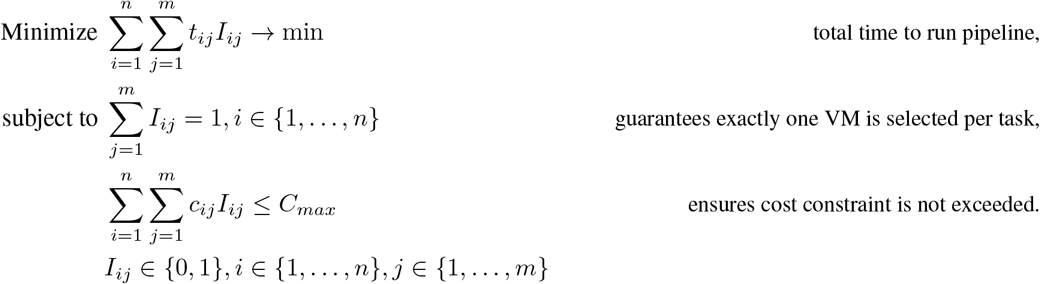

### 2.2 Distributed tasks

Sharding is a technique used to split a large task into smaller, more manageable pieces. The input data for the task is partitioned into shards based on a predetermined criterion, such as sequencing group or chromosome. Once the data is partitioned, it is distributed across multiple nodes for processing. Each node processes a subset of the data, which reduces the processing time and improves the overall performance of the system.

When a task is split into shards (as shown in Figure 2), we aggregate the shards and represent the task as a single vertex on the pipeline graph for several reasons. First, existing implementations of shard functionality don’t allow different virtual machines to be assigned to different shards, and each shard of the task is assigned the same type of virtual machine. Second, the goal of creating shards is usually to split the task into pieces as equally as possible. Therefore, we assume that the execution time and cost won’t be drastically different between shards. When collecting execution statistics, we define the cost to complete the distributed task as the sum of all shards’ costs, and the time to complete the distributed task as the maximum across shards’ run duration.

**Figure 2:**
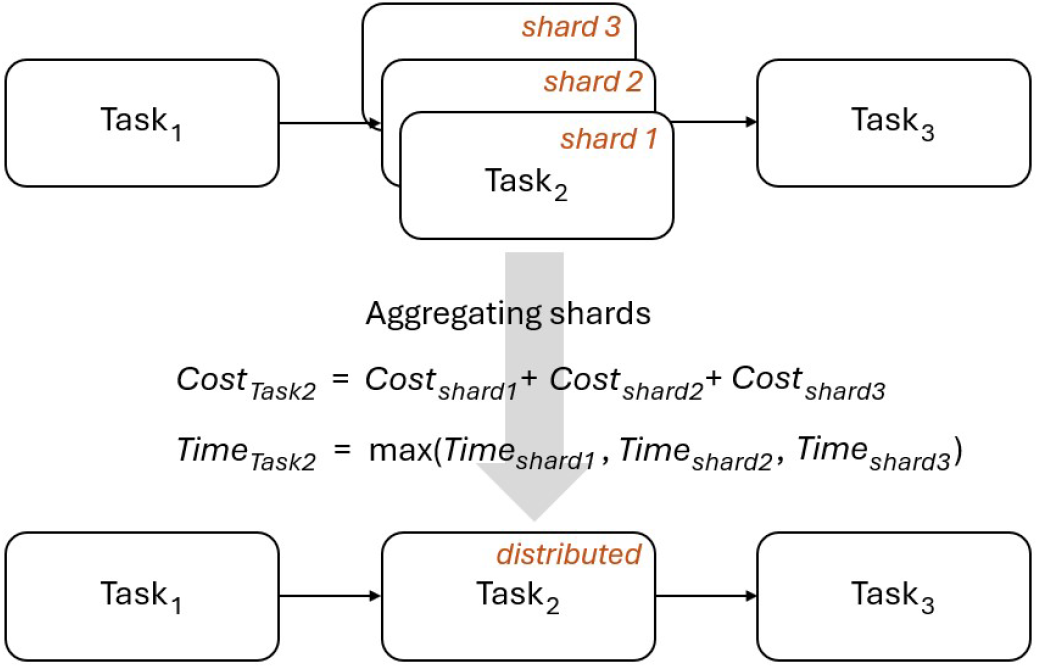
Example of distributed task and shards’ execution statistics aggregation

A pipeline with tasks split into shards can also be represented as a pipeline with non-linear topology (as shown in 2.3), it will increase the complexity of LP problem. It might be interesting to preform a comparison study to better understand the benefits from using non-linear topology versus the shard aggregation approach. This question is out of scope of this paper, and for simplicity, we will consider the distributed task as one vertex on pipeline graph.

### 2.3 Non-linear pipeline topology

The vast majority of genomics pipelines have a non-linear topology. In the non-linear case, the cost can still be computed as a sum of all costs, but the time to complete the pipeline should be computed differently. Let’s consider the pipeline presented in Figure 3. We can compute the time to complete it in the following way:

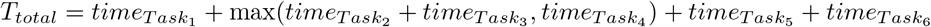

Unfortunately, we cannot replace the constraint in problem **LP 1** and the objective function in problem **LP 2** with the expression given above because it breaks the definition of an LP problem. However, there are tips and tricks that can help deal with non-linear functions, such as maximum, absolute value, etc., which frequently appear in real-world problems [13]. Usually, this is accomplished by creating additional variables.

**Figure 3:**
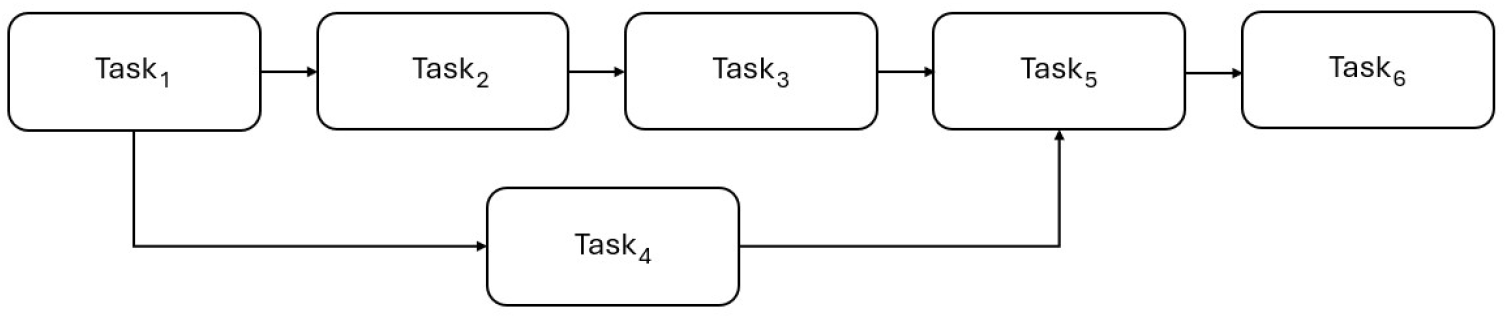
Example of non-linear pipeline topology

We introduce a variable *F*_*i*_ for each *Task*_*i*_, which describes amount of time required to complete a part of the pipeline from the start up to *Task*_*i*_. We also define a set of immediate predecessors for each vertex of the pipeline graph *ImPred*_*i*_. For example, for *Task*_5_ the set of immediate predecessors is *ImPred*_5_ = {*Task*_3_, *Task*_4_}, and two inequalities hold 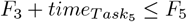 and 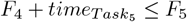. These inequalities ensure that *Task*_3_ and *Task*_4_ are completed prior to *Task*_5_, and then 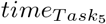 is required to complete the part of the pipeline from the start and up to *Task*_5_.

We also introduce *F*_*total*_, which describes the amount of time required to complete the whole pipeline from the start to the end. With additional variables, linear programming formulations for non-linear topology can be expressed as follows:

**LP 3 – Minimize cost subject to time constraint**

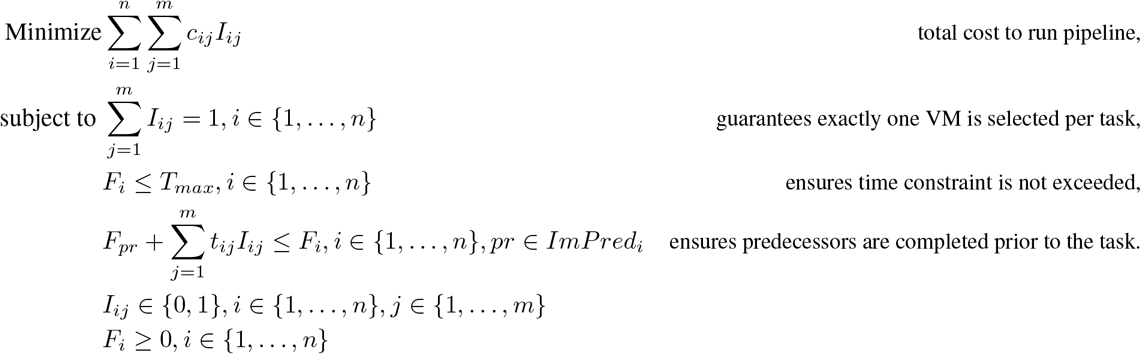

**LP 4 – Minimize time subject to cost constraint**

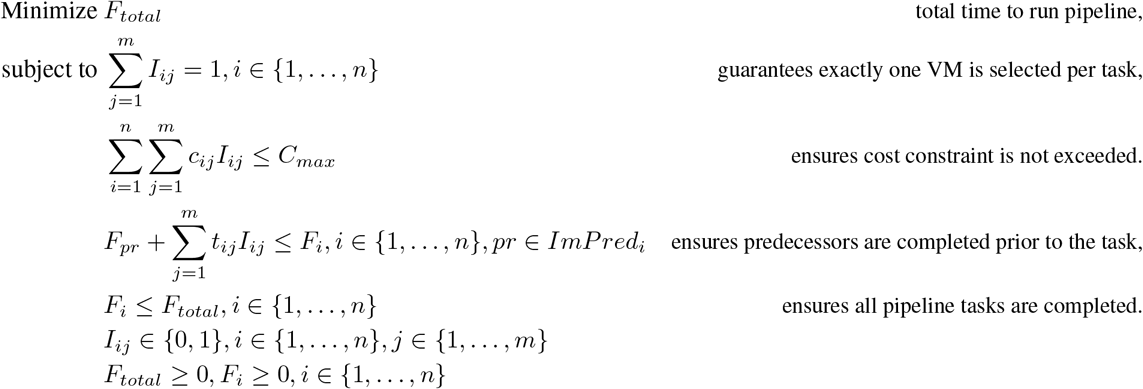

## 3 Application: optimizing genome data pre-processing pipeline

### 3.1 Pipeline and data

We leveraged LP problems formulated in the previous section to optimize UnmappedBamToAlignedBam workflow which implements the data pre-processing step of the Broad Institute Whole Genome Germline Single Sample Pipeline [15]. The version of the pipeline adapted to Azure is available in [16]. The UnmappedBamToAlignedBam workflow graph is presented in Figure 4: it has 13 tasks total, 5 of which are distributed. The list of immediate predecessors for each task is presented in Listing S4.

**Figure 4:**
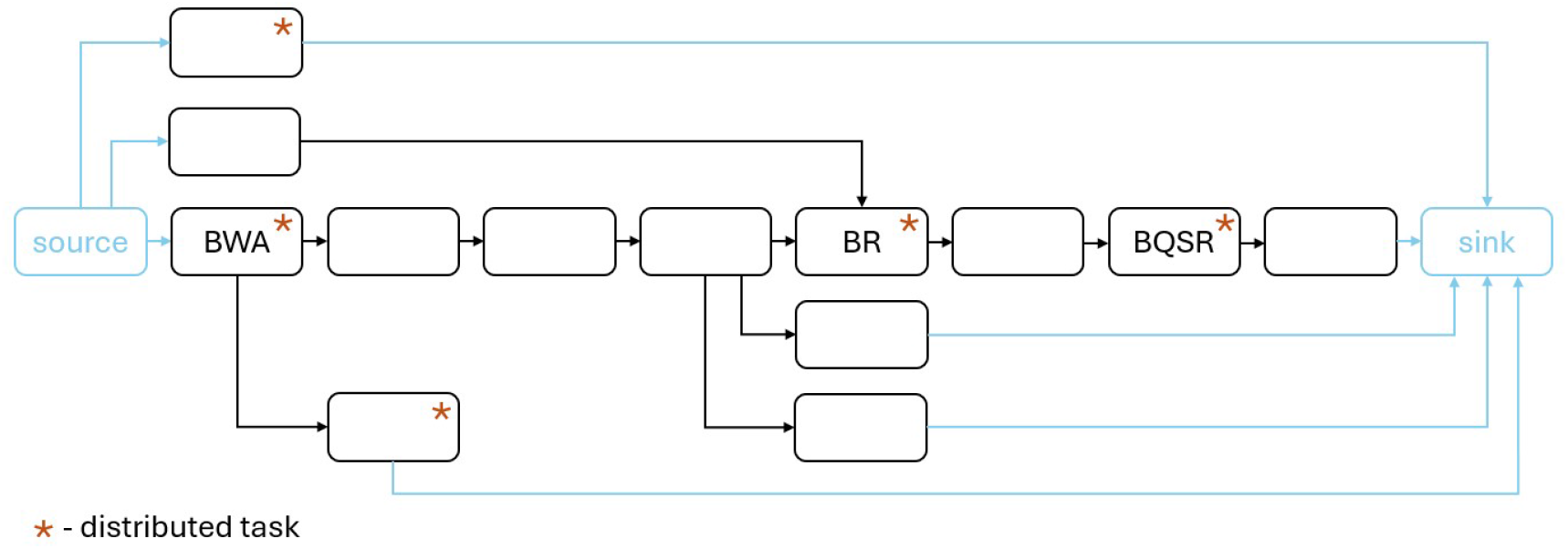
Pipeline graph for UnmappedBamToAlignedBam workflow. Tasks ‘sink’ and ‘source’ are introduced for implementation purposes to mark entry and exit points of the pipeline.

The pipeline employs different sharding strategies for distributed tasks. For SamToFastqAndBwaMemAndMba (labeled as BWA on the pipeline graph), CollectQualityYieldMetrics, and CollectUnsortedReadgroupBamQualityMetrics, one shard is created for each input file with no guarantees that the shards will be of equal size. For BaseRecalibrator and ApplyBQSR (labeled as BR and BQSR respectively), an attempt is made to split the input bam file into equal pieces with sequencing groupings. SamToFastqAndBwaMemAndMba is a compute-intensive task, and its performance significantly affects the performance of the entire pipeline. We use it to observe how the shard-per-file strategy affects pipeline performance in terms of cost and run duration.

We selected a subset of the 1000 Genomes Project dataset [17, 18]: 49 samples in total, each with 30x coverage. Of these, 46 samples have 12 unmapped bam files per sample, while the remaining 3 samples have 8, 13, and 14 files, respectively. Detailed sample statistics are presented in Table S1. To collect execution statistics, we randomly selected 5 samples out of the 46 samples with 12 files, as shown in Table 4. Genome assembly GRCh38 was used as a reference.

**Table 4:** Subset of samples used to collect pipeline execution statistics.

| Sample | Number of files | File size statistics (in GB) |  |  |  | Total size (in GB) |
| --- | --- | --- | --- | --- | --- | --- |
|  |  | mean | stdev | min | max |  |
| NA19017 | 12 | 4.13 | 0.19 | 3.95 | 4.45 | 49.6 |
| HG03436 | 12 | 3.77 | 1.17 | 2.88 | 5.41 | 45.19 |
| NA06984 | 12 | 3.64 | 0.36 | 3.13 | 4.04 | 43.64 |
| HG01982 | 12 | 3.62 | 0.68 | 2.95 | 4.65 | 43.45 |
| NA20764 | 12 | 3.6 | 0.44 | 2.85 | 3.92 | 43.14 |

### 3.2 Virtual machine assignments and execution statistics collection

We executed the UnmappedBamToAlignedBam workflow on Azure using a modified version of CromwellOn-Azure v3.0.0 [12]. The modifications were made to label each run with the sample and experiment name to simplify execution statistics collection and processing. No other functionality was changed.

Three experiments were defined: `default` VM assignment, `Dv2` family, and `Ddv4` family. By default, CromwellOn-Azure assigns the cheapest VM per hour that satisfies the task’s compute requirements, such as disk, memory, or number of CPUs. The list of defaults was saved and then passed as an additional input file for the pipeline run to guarantee the same defaults were used for all samples. For `Dv2` and `Ddv4` family experiments, we created VM assignment lists manually based on the tasks’ compute requirements. These lists were also passed as additional input files for the corresponding pipeline runs. All VM assignment lists are available in the Supplementary materials, see Listings S1–S3.

We were inspired by Intel’s benchmarking experiment [19] when choosing the VM families for our experiments. However, both Intel’s experiment and our experiments were conducted about two years ago. Therefore, the results presented in the next sections should not be considered as an optimal choice recommendation. The Azure VM offering has changed since then, with some families being deprecated and several new families being added. As a result, we highly recommend evaluating the currently available VM families to find up-to-date optimal compute resources.

To collect execution statistics we ran the pipeline three times per each sample defined in Table 4 and each experiment defined above, resulting in 45 runs in total. Then, for each task-experiment pair, we computed the average cost and execution time over runs and samples (see Table S2). The average cost and time to complete whole pipeline are presented in Table 5. We can observe that simple switch from `default` VM assignment to `Ddv4` family decreases time to complete the pipeline twice with a small increase in cost. We also computed the lower and upper bounds for cost and time that can be achieved by mixing VMs from the three experiments:

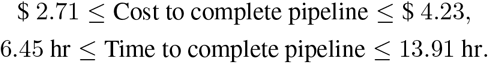

**Table 5:** Cost and time to complete pipeline for each experiment.

| Experiment | Cost (in \$) | Time (in hr) |
| --- | --- | --- |
| default | 3.26 | 13.88 |
| Dv2 | 3.55 | 8.23 |
| Ddv4 | 3.57 | 6.55 |

### 3.3 Implementing and solving LP problems

We used PuLP [20], a Python-based modeling language, to implement problems **LP 3** and **LP 4** (see Listings S5 and S6). PuLP can call a variety of open-source and commercial linear programming solvers, we used COIN-OR CBC [21], which is bundled with the PuLP package.

We selected **UC 2** from the two use cases described in the introduction, which involves minimizing the time to complete the pipeline with a restriction on cost. As shown above, the cost lower and upper bounds are $ 2.71 and $ 4.23 respectively. We took several points from the cost range [2.71, 4.23] and used each of them as the cost constraint *C*_*max*_, thus generating and solving several time minimization LP problems. The time matrix *T* and cost matrix *C* were obtained from the execution statistics presented in Table S2, pipeline graph was defined in JSON format (see Listing S4).

Table 6 displays the values used as cost constraints *C*_*max*_ and minimum times *T*_*min*_ to complete the pipeline corresponding to these constraints. *T*_*min*_ is the minimum of the objective function for problem **LP 4** with cost constraint *C*_*max*_. The results presented demonstrate the cost-time trade-off that we described in the introduction. For instance, if we are willing to spend $ 4.23 on each pipeline run, then the minimal time to complete the pipeline is 6.45 hours. On the other hand, if we do not want to spend more than $ 3.3 on each run, then we need a minimum of 6.53 hours. Table 6 also allows us to make a choice based on cost-time requirements specific to a research project. We decided to proceed with *C*_*max*_ = 2.71 and *T*_*min*_ = 6.82. Solution of LP problem with *C*_*max*_ = 2.71 is presented in Table 7.

**Table 6:** Cost constraints and corresponding minimal times to complete pipeline.

| $C_{max}$ (in \$) | $T_{min}$ (in hr) |
| --- | --- |
| 2.71 | 6.82 |
| 2.86 | 6.53 |
| 3.01 | 6.53 |
| 3.17 | 6.53 |
| 3.32 | 6.53 |
| 3.47 | 6.45 |
| 3.62 | 6.45 |
| 3.77 | 6.45 |
| 3.93 | 6.45 |
| 4.08 | 6.45 |
| 4.23 | 6.45 |

**Table 7:** Recommended VM assignment when minimizing time with cost constraint *C*_*max*_ = 2.71.

| Task | Recommended experiment | Recommended VM |
| --- | --- | --- |
| ApplyBQSR | Dv2 | Standard_D3_v2 |
| BaseRecalibrator | Ddv4 | Standard_D4d_v4 |
| CheckContamination | Ddv4 | Standard_D4d_v4 |
| CollectQualityYieldMetrics | Ddv4 | Standard_D2d_v4 |
| CollectUnsortedReadgroupBamQualityMetrics | Ddv4 | Standard_D2d_v4 |
| CreateSequenceGroupingTSV | Ddv4 | Standard_D2d_v4 |
| CrossCheckFingerprints | Ddv4 | Standard_D2d_v4 |
| GatherBamFiles | Ddv4 | Standard_D4d_v4 |
| GatherBqsrReports | Ddv4 | Standard_D2d_v4 |
| MarkDuplicates | Ddv4 | Standard_D8d_v4 |
| SamToFastqAndBwaMemAndMba | default | Standard_F16s_v2 |
| SortSampleBam | Ddv4 | Standard_D8d_v4 |
| SumFloats | Ddv4 | Standard_D2d_v4 |

### 3.4 Using recommended VM assignment to process full set of samples

We analyzed the full set of 49 samples using the compute resources recommended in Table 7. One run was performed for each sample, and for the majority of samples, the pipeline run was completed within (or very close to) the cost-time boundaries defined by *Cost* ≤ $ 2.71 and *Time* ≤ 6.82 hr (see the bottom left section of Figure 5). Sample NA12872 is not present on the figure, as the run for this sample failed due to its large size and lack of disk space for processing a file by one of the non-distributed tasks. Samples that appear on the right section of Figure 5 are characterized by a larger standard deviation of sample file sizes, a large total sample size, or a combination of both (see Table 8).

**Table 8:** Top 15 samples by size; for samples in **bold** cost or time to complete pipeline exceeded limits *Cost* ≤ $ 2.71 and *Time* ≤ 6.82 hr.

|  | Sample | Number of files | File size statistics (in GB) |  |  |  | Total size (in GB) |
| --- | --- | --- | --- | --- | --- | --- | --- |
|  |  |  | mean | stdev | min | max |  |
| 1 | **NA12872** | 12 | 4.6 | 0.97 | 3.21 | 5.56 | **55.25** |
| 2 | **HG03006** | 12 | 4.34 | **2.14** | 2.13 | 7.2 | **52.07** |
| 3 | **HG01565** | 12 | 4.32 | 0.27 | 3.85 | 4.63 | **51.81** |
| 4 | **HG02855** | 12 | 4.19 | 0.44 | 3.66 | 4.8 | **50.25** |
| 5 | **NA19017** | 12 | 4.13 | 0.19 | 3.95 | 4.45 | **49.6** |
| 6 | **NA20869** | 12 | 4.05 | **2.05** | 2.43 | 6.89 | 48.6 |
| 7 | NA19035 | 12 | 3.89 | 0.58 | 3.13 | 4.67 | 46.73 |
| 8 | HG03727 | 12 | 3.85 | 0.14 | 3.66 | 4.15 | 46.24 |
| 9 | **NA12340** | 14 | 3.3 | **3.21** | 1.93 | 11.65 | 46.15 |
| 10 | HG01583 | 12 | 3.84 | 0.15 | 3.67 | 4.08 | 46.08 |
| 11 | HG03742 | 12 | 3.81 | 0.23 | 3.57 | 4.16 | 45.71 |
| 12 | NA20502 | 12 | 3.77 | 0.45 | 3.35 | 4.42 | 45.3 |
| 13 | NA20509 | 12 | 3.77 | 0.15 | 3.57 | 4.03 | 45.24 |
| 14 | NA18995 | 12 | 3.77 | 0.11 | 3.61 | 3.91 | 45.22 |
| 15 | **HG03436** | 12 | 3.77 | **1.17** | 2.88 | 5.41 | 45.19 |

**Figure 5:**
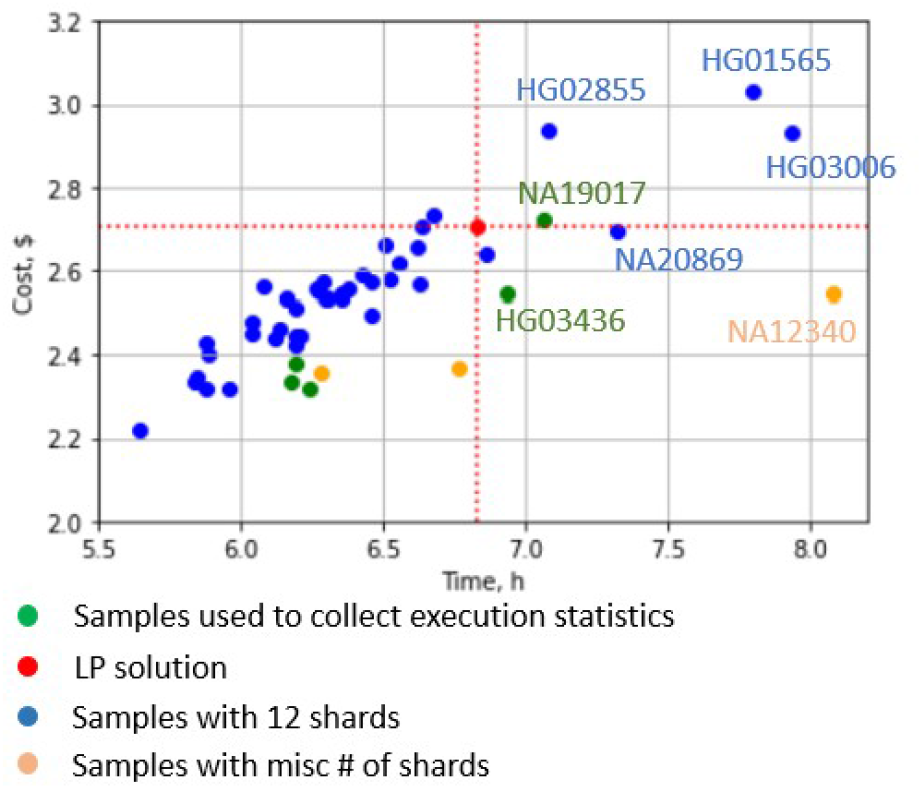
Cost and time to complete data pre-processing pipeline with recommended VM assignment, each point corresponds to one sample.

Samples HG03436, NA20869, and NA12340 illustrate how the duration of the pipeline run is affected by the variability in sample file sizes. As previously mentioned, the task SamToFastqAndBwaMemAndMba is a distributed task where one shard is created for each input file, and the same type of VM is assigned to all shards of the task. For sample NA12340, input files of size 1.93 GB and 11.65 GB are processed in parallel on two VMs of the same type (Standard_F16s_v2). Since larger files require more processing time, variability in file sizes leads to variability in shard processing times. The task SamToFastqAndBwaMemAndMba is completed when the longest shard is completed, and for NA12340, it takes longer compared to evenly distributed samples. At the same time, the total amount of processed data for NA12340 (46.15 GB) is similar to that of other samples, so we do not observe an increase in the cost of the pipeline run. Overall, the uneven distribution of samples across input files shifts samples to the right on the Time vs. Cost graph, whilst an increase in total sample size affects both cost and time to complete the pipeline, and shifts samples to the top right (see sample HG01565).

## 4 Discussion

While our assumptions such as average execution statistics, aggregated shards, and multiple-choice problem formulation help simplify the LP problem and make it more accessible to non-experts, they also introduce limitations to the approach described above. In this section, we discuss several enhancements that can be implemented and the scenarios in which they might be useful.

### Dealing with distributed tasks

As demonstrated earlier, combining shards into a single node on the pipeline graph may not be effective when there is an uneven distribution of inputs across shards. If this scenario is anticipated, an alternative approach to consider is to represent each shard with a separate node and use a non-linear pipeline topology.

### Cloud compute resources are limited

We made the assumption that VMs used for experiments are available in any quantity when we wrote multiple-choice problems and formulated LP problems. This assumption might be acceptable if samples are analyzed in small quantities in sequential order. For instance, if a few new samples come every 8 hours, they’re sent for analysis right away and it’s completed before the next batch comes. In this scenario, resource capacity issues might be prevented with careful evaluation of compute requirements, Cloud quotas and competing workloads. If the capacity limits for some virtual machines are reached when analyzing one sample and changing the quota is not possible, then a capacity constraint must be added to the LP problems.

When analyzing samples at a large scale, capacity limits may be easily reached by multiple distributed tasks with hundreds or thousands of shards running in parallel. For instance, one HaplotypeCaller task from Whole Genome Germline Single Sample Pipeline requires 50 shards for a 30x coverage sample. In this case, it might be beneficial to optimize resource utilization in general and consider a resource allocation and scheduling problems that are well-known in Cloud computing. We recommend reviewing the current state of research in this field and carefully evaluating the complexity of the solution that your project requires.

### Is execution statistics required?

We assumed that the researcher is willing to collect execution statistics before optimizing the pipeline, which might be impossible or cost-prohibitive in some cases. Additionally, we used average cost and execution time values to solve LP problems, which resulted in the loss of information about individual sample size, task cost, and execution time distributions. We also did not take into account the input size of incoming sample to process.

If we can accurately predict task execution time and cost for each sample, then VM assignment recommendation can be done individually as well. When historical execution data is available, various machine learning techniques can be used to predict task runtime for new samples based on their size, workflow structure, etc. [22, 23, 24, 25, 26] However, approaches relying on historical data cannot be applied to workflows without any historical traces or workflows with changed parameters and configurations on any kind of cluster. In such cases, methods employing microbenchmarks for predicting the runtimes of tasks before they are executed might be very useful [27].

## 5 Conclusion

In this paper, we present a linear programming method that can help bioinformatics researchers optimize the execution of scientific pipelines for either cost or time. Our method is designed to work with all scientific pipelines that have a directed acyclic graph topology. The implementation we present uses execution data from representative samples to determine the optimal compute needed for the pipeline execution. The same methodology might be used with individual task runtimes predicted on historical data or microbenchmarks. In the future, we want to explore the limitation of compute resources and capacity when dealing with competing interests, such as running competing pipelines or scaling a single pipeline at thousands of parallel samples.

## Supplementary materials

### Compute resources used for experiments

**Listing S1:**
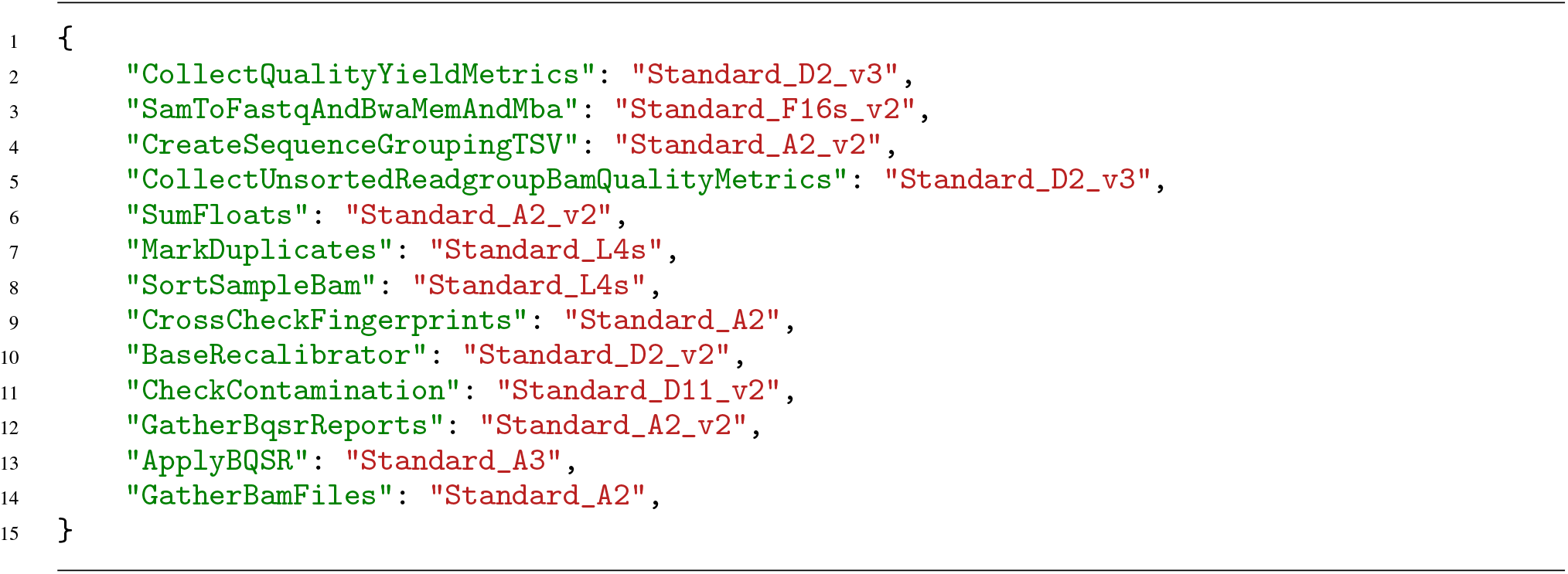
Default virtual machine assignment made by CromwellOnAzure

**Listing S2:**
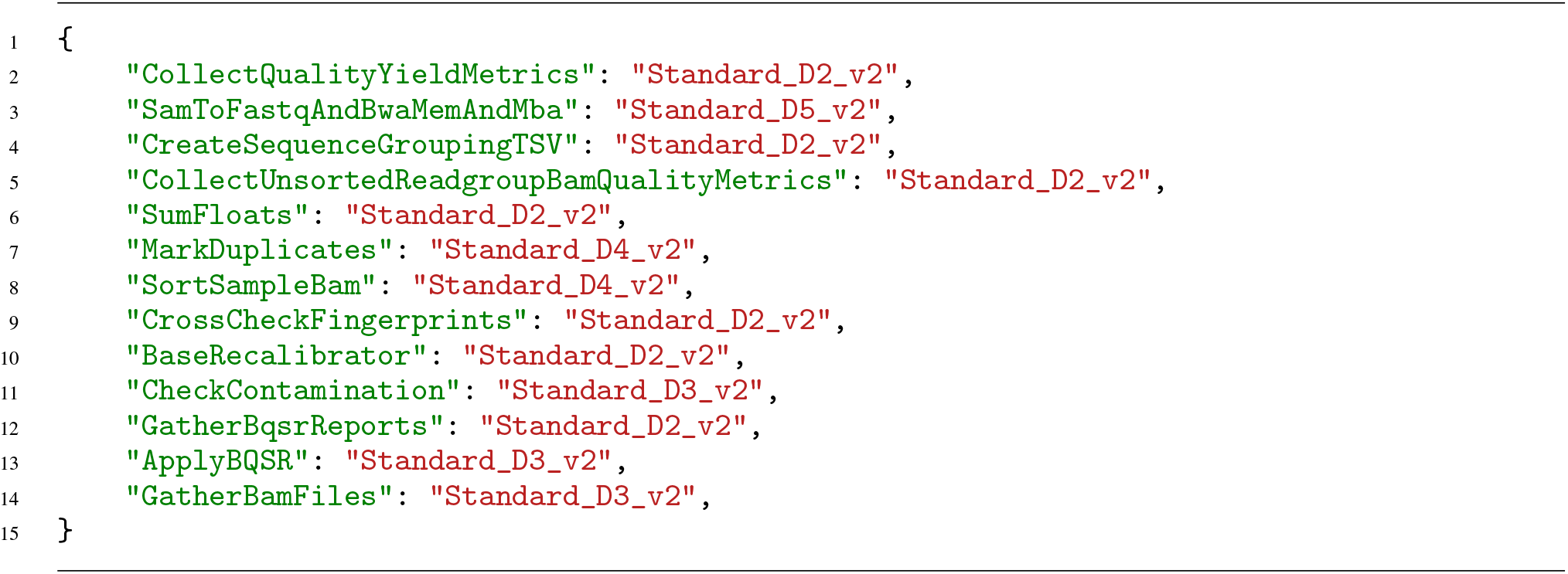
Virtual machine assignment for Dv2 family experiment

**Listing S3:**
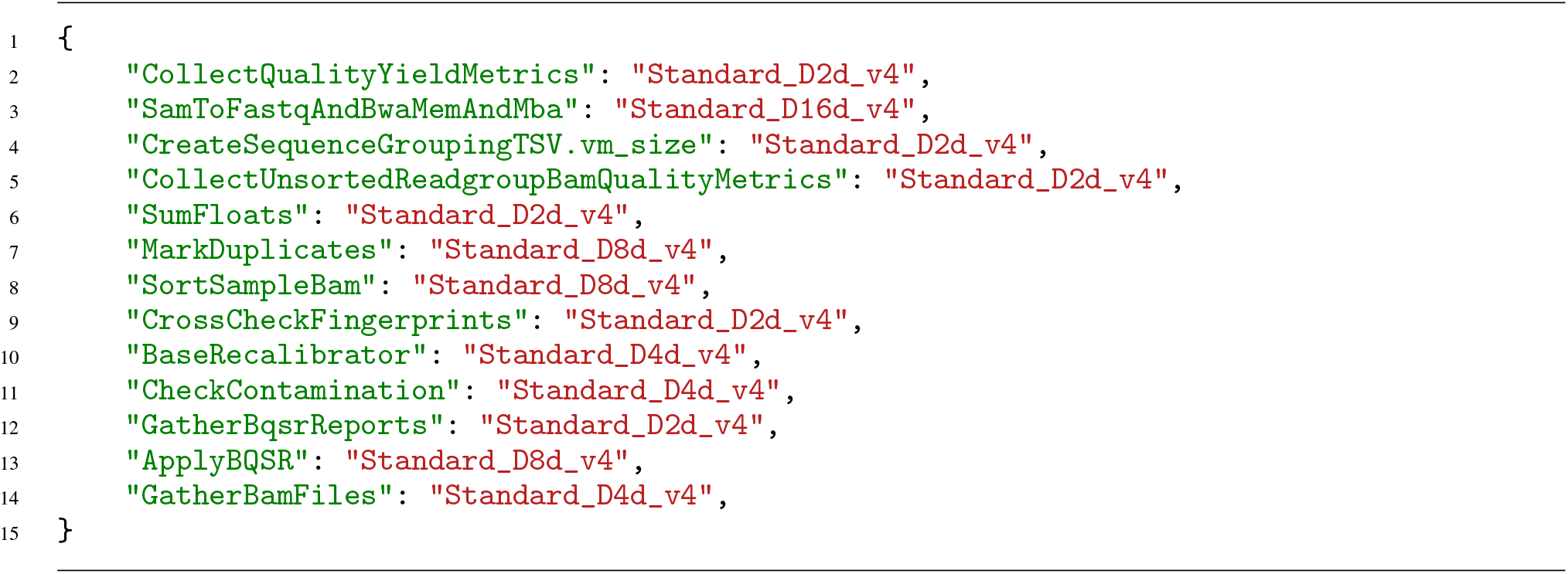
Virtual machine assignment for Ddv4 family experiment

### UnmappedBamToAlignedBam workflow graph

**Listing S4:**
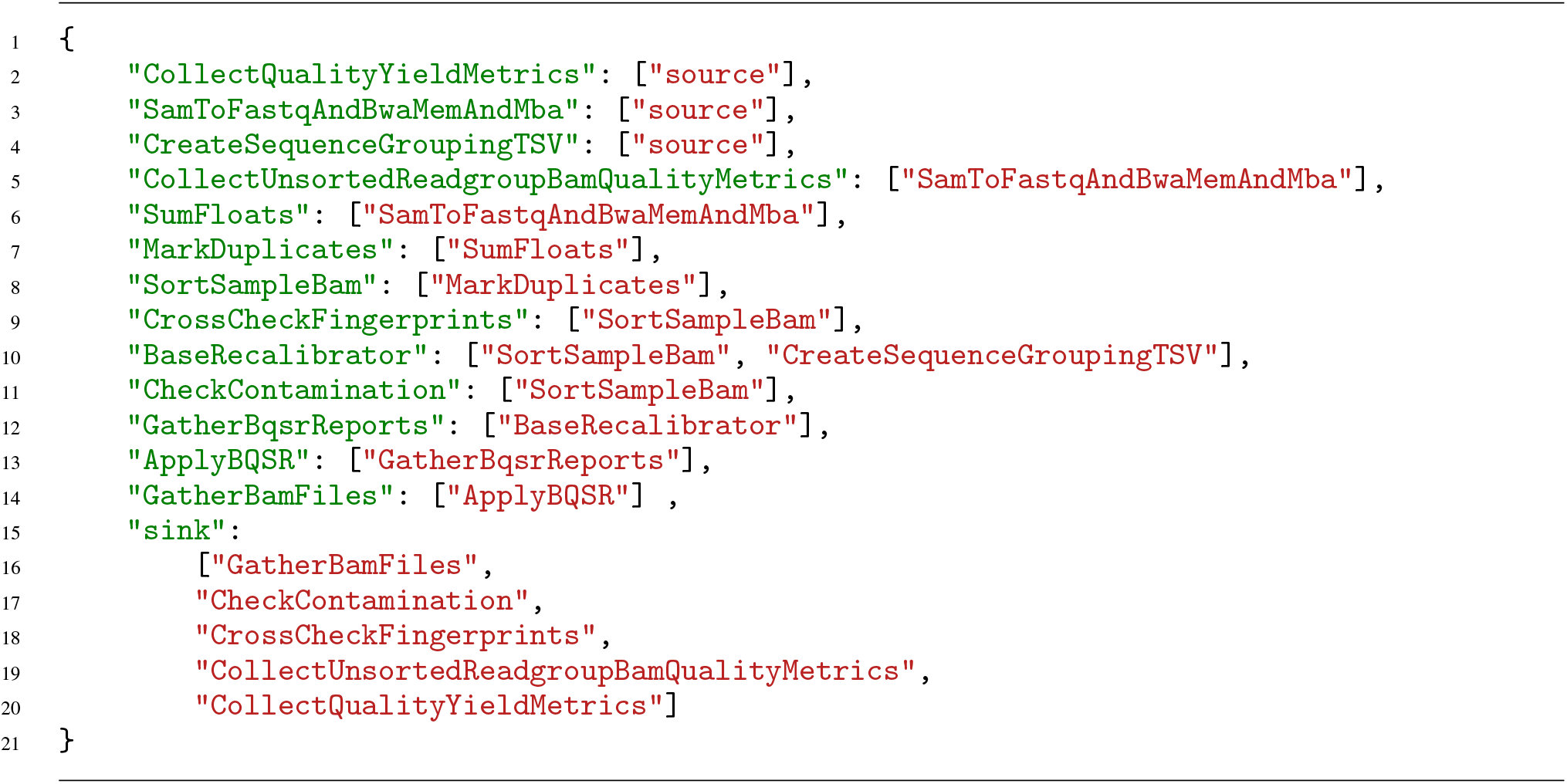
Default virtual machine assignment made by CromwellOnAzure purposes to mark entry and exit points of the pipeline.

### Implementation of LP problems for non-linear pipeline topology using PuLP

Below we assume that `pipeline` is an instance of a to-be-implemented `Pipeline` class with the following properties:

**Listing S5:**
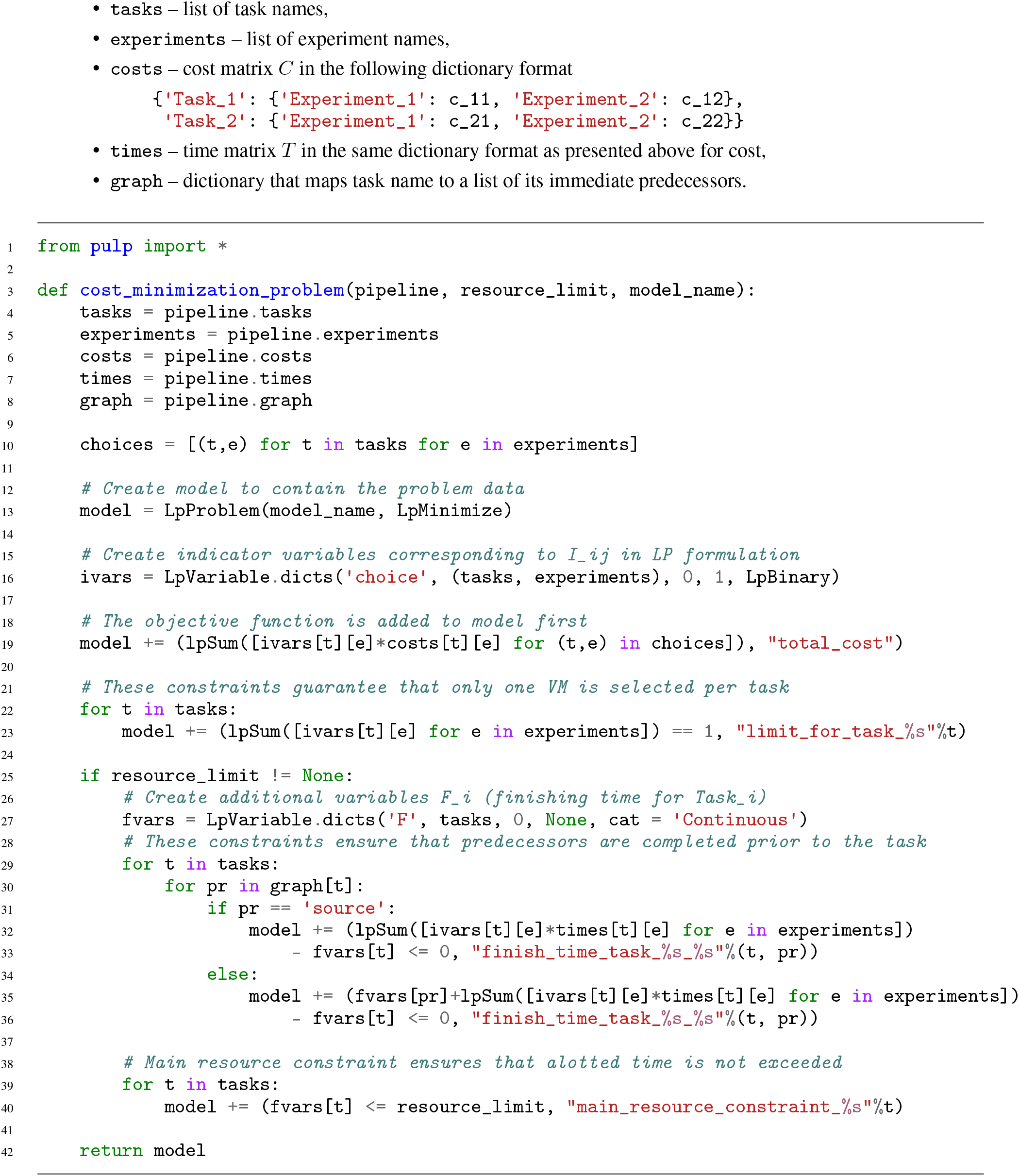
PuLP model for cost minimization problem

**Listing S6:**
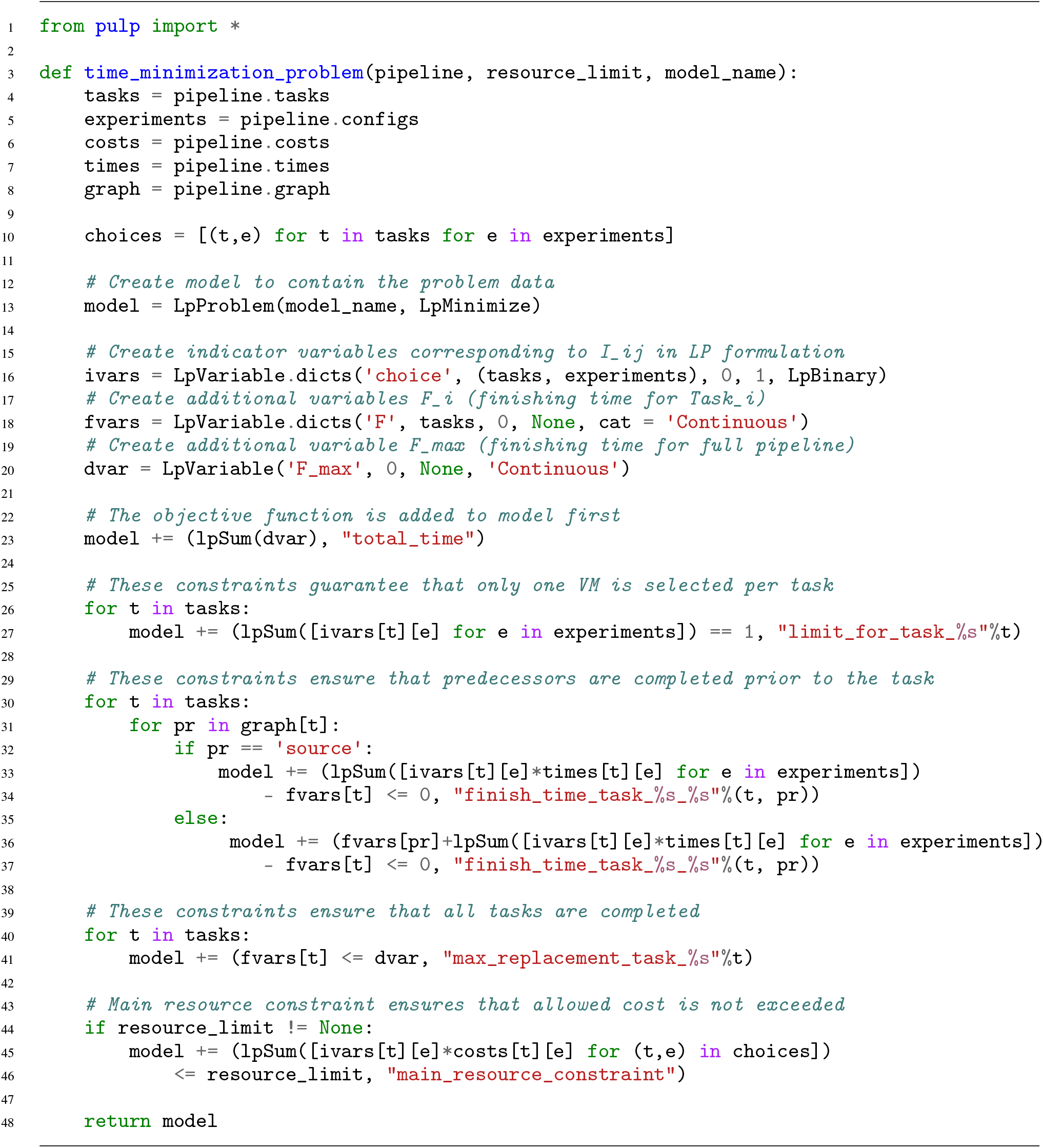
PuLP model for time minimization problem

### List of samples and execution statistics

**Table S1:**
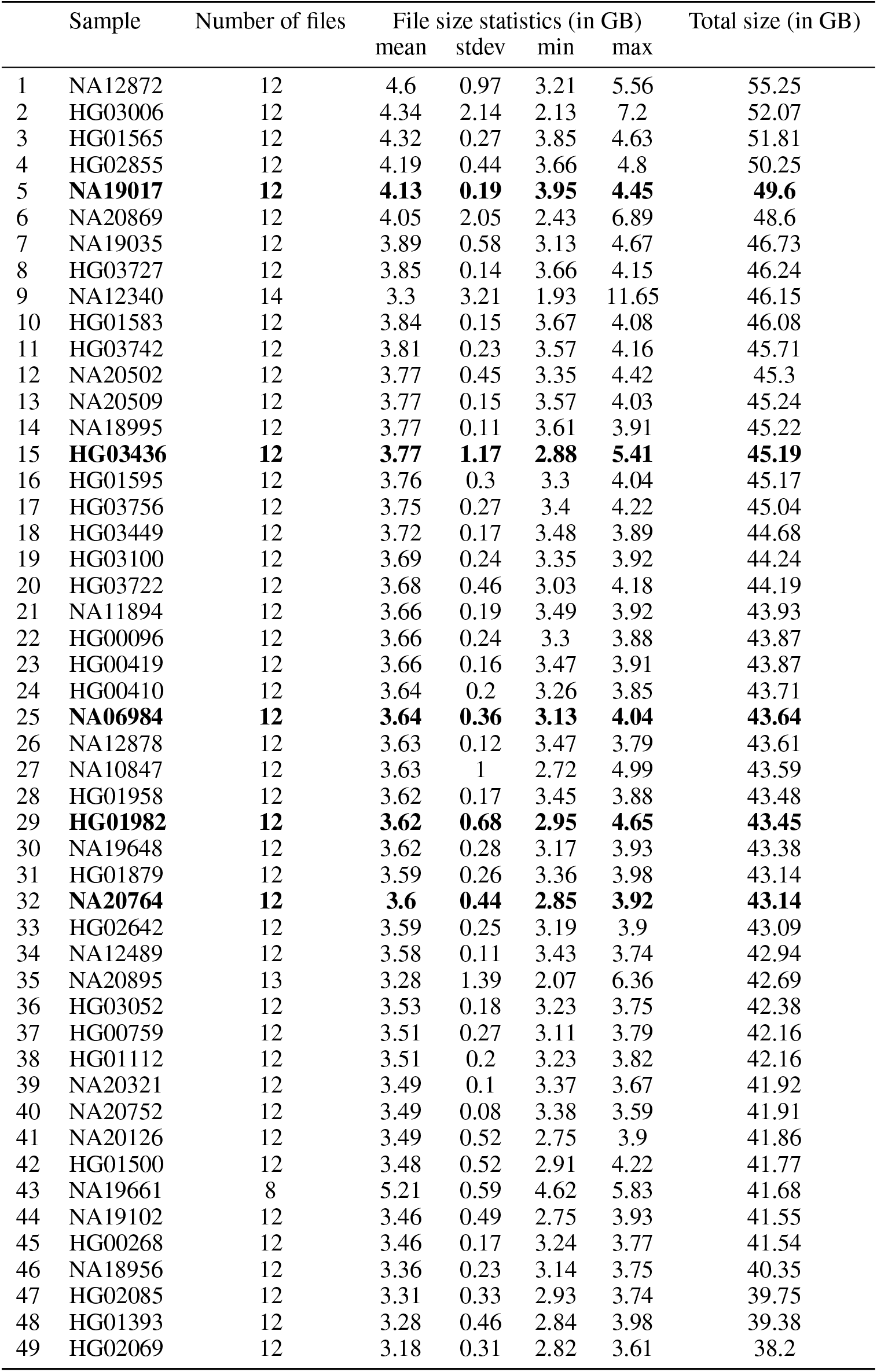
Full list of samples analyzed by data pre-processing pipeline, ordered by total sample size. Samples in **bold** were used to collect pipeline execution statistics presented in Table S2

**Table S2:** Pipeline execution statistics

| Task | Experiment | Cost (in \$) | Time (in hr) |
| --- | --- | --- | --- |
| ApplyBQSR | Ddv4 | 0.591 | 0.459 |
| ApplyBQSR | Dv2 | 0.524 | 0.757 |
| ApplyBQSR | default | 0.903 | 1.39 |
| BaseRecalibrator | Ddv4 | 0.338 | 0.531 |
| BaseRecalibrator | Dv2 | 0.356 | 1.19 |
| BaseRecalibrator | default | 0.37 | 1.19 |
| CheckContamination | Ddv4 | 0.0302 | 0.668 |
| CheckContamination | Dv2 | 0.043 | 0.769 |
| CheckContamination | default | 0.0425 | 1.15 |
| CollectQualityYieldMetrics | Ddv4 | 0.0175 | 0.0824 |
| CollectQualityYieldMetrics | Dv2 | 0.0283 | 0.132 |
| CollectQualityYieldMetrics | default | 0.0227 | 0.19 |
| CollectUnsortedReadgroupBamQualityMetrics | Ddv4 | 0.0381 | 0.175 |
| CollectUnsortedReadgroupBamQualityMetrics | Dv2 | 0.0615 | 0.248 |
| CollectUnsortedReadgroupBamQualityMetrics | default | 0.0503 | 0.354 |
| CreateSequenceGroupingTSV | Ddv4 | 0.000223 | 0.00986 |
| CreateSequenceGroupingTSV | Dv2 | 0.000314 | 0.0108 |
| CreateSequenceGroupingTSV | default | 0.000358 | 0.0199 |
| CrossCheckFingerprints | Ddv4 | 0.00509 | 0.225 |
| CrossCheckFingerprints | Dv2 | 0.00858 | 0.296 |
| CrossCheckFingerprints | default | 0.0126 | 0.523 |
| GatherBamFiles | Ddv4 | 0.0272 | 0.602 |
| GatherBamFiles | Dv2 | 0.0368 | 0.659 |
| GatherBamFiles | default | 0.117 | 4.86 |
| GatherBqsrReports | Ddv4 | 0.000737 | 0.0326 |
| GatherBqsrReports | Dv2 | 0.00133 | 0.0459 |
| GatherBqsrReports | default | 0.0022 | 0.122 |
| MarkDuplicates | Ddv4 | 0.194 | 2.15 |
| MarkDuplicates | Dv2 | 0.28 | 2.5 |
| MarkDuplicates | default | 0.202 | 2.93 |
| SamToFastqAndBwaMemAndMba | Ddv4 | 2.17 | 1.03 |
| SamToFastqAndBwaMemAndMba | Dv2 | 1.97 | 0.931 |
| SamToFastqAndBwaMemAndMba | default | 1.37 | 1.01 |
| SortSampleBam | Ddv4 | 0.157 | 1.74 |
| SortSampleBam | Dv2 | 0.24 | 2.14 |
| SortSampleBam | default | 0.163 | 2.37 |
| SumFloats | Ddv4 | 0.000225 | 0.00998 |
| SumFloats | Dv2 | 0.000261 | 0.009 |
| SumFloats | default | 0.000459 | 0.0255 |

